# Approaches to generating virtual patient cohorts with applications in oncology

**DOI:** 10.1101/2022.05.24.493265

**Authors:** Anudeep Surendran, Justin Le Sauteur-Robitaille, Dana Kleimeier, Jana Gevertz, Kathleen Wilkie, Adrianne L. Jenner, Morgan Craig

## Abstract

Virtual clinical trials (VCTs) have gained popularity for their ability to rationalize the drug development process using mathematical and computational modelling, and to provide key insights into the mechanisms regulating patient responses to treatment. In this chapter, we cover approaches for generating virtual cohorts with applications in cancer biology and treatment. VCTs are an effective tool for predicting clinical responses to novel therapeutics and establishing effective treatment strategies. These VCTs allow us to capture inter-individual variability (IIV) which can lead to diversity in patient drug responses. Here we discuss three main methodologies for capturing IIV with a VCT. First, we highlight the use of population pharmacokinetic (PopPK) models, which extrapolate from empirical data population PK parameters that best fits the individual variability seen in drug disposition using non-linear mixed effects models. Next, we show how virtual patients may be sampled from a normal distribution with mean and standard deviation informed from experimental data to estimate parameters in a mechanistic model that regulates drug PKs. Lastly, we show how optimization techniques can be used to calibrate virtual patient parameter values and generate the VCT. Throughout, we compare and contrast these methods to provide a broader view of the generation of virtual patients, and to aid the decision-making process for those looking to leverage virtual clinical trials in their research.

## INTRODUCTION

Cancer is a heterogeneous disease with complex (sub)types, genetic compositions, and tumour spatial arrangements, all of which make designing and scheduling effective and minimally toxic cancer treatments more challenging. Despite the long-term concerted investment in highly intensive cancer research, the goal of precision and personalized medicine remains largely unrealized. The difficulty establishing new cancer treatment strategies is compounded by the complexity of the drug development pipeline, since getting a drug to market is a lengthy and expensive process. There have been many recent encouraging advances in cancer therapy, particularly the development of immunotherapies like T-VEC (an oncolytic virus to treat late-stage melanoma) (Andtbacka et al. 2015, 2016), and immune checkpoint inhibitors including nivolumab and pembrolizumab (Seidel et al. 2018). However, a number of disappointing trial results have highlighted the need for improved predictive and quantitative models to help guide clinical trials in oncology. Quantitative systems pharmacology (QSP) seeks to answer this call by developing systemic mathematical and computational models to explore dosing ranges and therapeutic regimens prior and concurrent to clinical trials.

Drug development in oncology and in general relies on the use of clinical trials. A randomized clinical trial evaluates new medical approaches by randomly dividing participants into separate groups, or arms. In these trials, either a new medical approach is compared to a placebo, or to an existing treatment (e.g. standard drug) in a non-inferiority trial. In a cross-over trial, multiple study arms may receive both treatments after a washout period which is calculated according to each drug’s half-life (Brown 1980; Piantadosi 2017). In all scenarios, the trial population is randomly separated into study arms to satisfy the requirement of equally distributed cohorts for reproducibility and comparability (Bland and Altman 2011). This randomization is of course never perfect or identical, something which the virtual clinical trials we will describe later aim to address. In early phases, drug tolerability is tested and dose escalation is performed before a drug’s efficacy in treating the target is measured (Lipsky and Sharp). At each stage, between patient variability (e.g. genetic, physiological etc.) leads to diversity in patient drug responses (Alfonso et al. 2020). Accounting for this inherent variability is often a significant obstacle for establishing effective and tolerable treatment schedules. Improperly estimating heterogeneity when planning trials, or observing a high degree of interindividual variability (IIV) contributes to decreased drug development success rates, which explains the high degree of attrition along the drug development pipeline (Kozłowska et al. 2019). In response, new approaches encompassing virtual clinical trials (VCT) have been increasingly integrated to (pre-)clinical drug development efforts as a means to quantify the effects of variable environmental, spatial, and genetic etc. factors on therapeutic regimens and patients (Alfonso et al. 2020).

VCTs arose with the emergence of QSP (Polasek and Rostami-Hodjegan 2020; Ma et al. 2021) and enable finding potentially non-intuitive drug regimens that help to increase drug approval success rates and, in turn, reduce drug costs (Alfonso et al. 2020). To that end, VCTs have been shown to be an effective tool in mitigating various challenges arising at different stages of the drug development pipeline, particularly understanding heterogeneous responses to novel therapeutics, establishing efficacious treatment strategies and treatment schedules, and precision dosing in individual patients (Polasek and Rostami-Hodjegan 2020). Virtual clinical trials have been applied to a broad range of diseases including HIV (Stadeli and Richman 2013; Kirtane et al. 2018), tuberculosis (Pitcher et al. 2018), SARS-COV-2 (Jenner et al. 2021a), sepsis (Clermont et al. 2004; An 2004; Vodovotz and Billiar 2013), diabetes (Visentin et al. 2014; Gyuk et al. 2019), cardiovascular diseases (Corral-Acero et al. 2020), and different tumours (including breast (Switchenko et al. 2019; Corral-Acero et al. 2020), brain (Agosti et al. 2018), melanoma (Barish et al. 2017; Cassidy and Craig 2019; Milberg et al. 2019), and lung (Jafarnejad et al. 2019; Sayama et al. 2021)). The successful implementation of VCTs heavily depends on the ability to generate heterogeneous virtual patients that mimic a broad spectrum of patients with a multitude of disease presentations, as would be observed in real-world clinical studies.

As detailed in later sections, VCTs generally follow the same basic steps. First, a mathematical or computational model of a disease is constructed using prior information and domain expertise. Model parameters are estimated from existing biological studies, targeted experimentation, or ongoing or completed trials. Next, the sensitivity of model predictions to perturbations in parameter values is determined through local or global sensitivity analyses. To simulate a population-level response, with variations in individual patient responses (Cassidy and Craig 2019), the mathematical model is solved for each individual-based parameter set. These patient-mimicking parameter sets are informed by the sensitivity analyses and constructed either statistically, by imposing physiologically-appropriate ranges and distributions on the values, or by probabilistic data-fitting, to ensure that model predictions lie within ranges of experimental or clinical observations, using optimization schemes. Importantly, parameter value ranges may be pruned to ensure that model predictions recapitulate physiologically reasonable ranges and observations (Allen et al. 2016). If generated statistically, virtual patients are selected by randomly sampling each parameter value from the chosen distributions, and the resulting parameter set is accepted into the trial if predicted outcomes are within acceptable deviations from the known outcomes. If generated through probabilistic data-fitting, virtual patients are constructed as a set of parameter values by each successful fit of the optimization algorithm, and are thus automatically accepted into the trial. However generated, virtual patients accepted into the VCT are twinned/cloned and assigned to multiple identical cohorts. Therapeutic outcomes for each virtual patient in each cohort are then simulated and compared using a variety of statistical techniques. This virtual study design is evocative of a crossover study, with the advantage that patients can be assigned to multiple cohorts at once, implying that differences observed between cohorts can be attributed to mechanistic causes (as cohorts are identical). VCTs using the strategies described above have been applied to non-chronic diseases (Alfonso et al. 2020; Jenner et al. 2021a) and to identify causal mechanisms controlling outcomes (Cassidy and Craig 2019; Alfonso et al. 2020; Jenner et al. 2021a).

Typically, for large models with many parameter values, not all parameters will be assigned to the patient-specific parameter sets, as leaving some parameters constant across the population can simplify the VCT construction and analysis. When choosing which model parameters to fix as constants for the whole population, and which to vary across virtual patients, there are three key aspects of the mathematical model to consider: identifiability, sensitivity, and the biological interpretation of parameters. Model identifiability refers to the ability of model parameter values to be uniquely determined by comparing to observations. For example, if data is limited or incomplete, certain parameter values may not be well constrained by data fitting algorithms, resulting in a wide range of acceptable values that qualitatively fit the data. Performance may then be improved by identifying these parameters first and estimating their values from other sources or studies if possible. Or, these parameters may represent significant biological mechanisms that are a desirable addition to the patient-specific parameter set. Model sensitivity analysis compares the dependence of model predictions to small changes in each parameter value. A parameter with high sensitivity coefficient, for example, may be useful to capture population-level variance with minimal dimensionality in the patient-specific parameter sets. However, a parameter with low sensitivity coefficient may represent a significant biological mechanism that is desired in the variable parameter set. Thus, the decision of which parameters to fix, and which to allow to vary involves integration of all three above-mentioned key aspects. Once the composition of the variable parameter set is determined, these parameters may be used as potential biomarkers (Jafarnejad et al. 2019) to classify the virtual population.

For a number of reasons, VCTs are particularly attractive in cancer therapy development. Recent estimates put the probability that a candidate drug entering Phase I study today will obtain regulatory approval at around 10% (Hay et al. 2014a), requiring around $3.953 billion USD for out-of-pocket and capitalized research and development costs (DiMasi et al. 2016) over 13 years (Mohs and Greig 2017). Yet, this probability is even lower in oncology (Hay et al. 2014b), demonstrating the need for improved approaches in this area. Among the therapies that have met with disappointing real-world trial results are new cancer therapy strategies that aim to use the body’s natural immune response against tumours. One reason for disappointing trial results with such immunotherapies is the complex, heterogeneous, and dichotomous nature of tumour-immune interactions (Wilkie and Hahnfeldt 2017; Jafarnejad et al. 2019; Wang et al. 2020). Another reason for the high failure rate in cancer clinical trials is the evolution of drug resistance (Bozic et al. 2010; Tirosh et al. 2016; Craig et al. 2019).

Drug resistance in particular can be addressed using VCTs, as in Emilia Kozłowska et al. 2018 where the authors constructed a model to find promising drug regimens to prevent platinum resistance of ovarian tumours typically treated with surgery and platinum-based chemotherapy. Unfortunately, relapses of ovarian tumours are highly frequent, but combination therapy with different platinum-taxanes (paclitaxel or docetaxel) can increase the amount of platinum sensitive cancer cells and the time to tumour relapse by administering drugs in six different combinations, optimally with three-to-four drugs (Emilia Kozłowska et al. 2018). A second study by Kozłowska et al. compared treatments using three different drugs^9^. A combination of trientine, a copper chelating agent, and birinapant in biomarker-selected treatments, was compared with a biomarker-unselected treatment with wWee1 inhibitor resulting in an increased survival of Wee1 inhibitor-treated virtual patients. Other such VCTs have also been investigated in the context of preventing the evolution of resistance^9,39–41^.

To further improve cancer treatment, VCTs have focused on the reduction of toxicity (and therefore increased tolerance) to improve therapeutic outcomes, and have studied new immunotherapies (Barish et al. 2017; Sové et al. 2020). For example, QSP-IO is a platform for modeling immuno-oncology (IO), which accommodates varying degrees of model complexity based on specific research questions (Sové et al. 2020). To build QSP-IO, the authors implemented several different modules, including aspects of T cell behaviour or the effects of immune checkpoint inhibitors (Jafarnejad et al. 2019; Ma et al. 2021). By applying those platforms on data from triple-negative breast cancer clinical trials with administration of placebo and nab-paclitaxel or atezolizumab and nab-paclitaxel concurrent therapies of atezolizumab and nab-paclitaxel were found to be the best therapy options (Wang et al. 2021). Similarly, VEPART (Barish et al. 2017) is a tool for identifying robust optimal treatment protocols integrating experimental data, mathematical modeling, and statistical analyses that was applied to a melanoma mouse model treated with immunostimulatory oncolytic viruses and dendritic cell vaccines to investigate optimal dose scheduling. Barish et al. 2017 found that, subject to a number of constraints on dose and treatment length, only one optimal combination (three days of oncolytic virus, followed by three days of dendritic cell therapy) led to total tumor eradication in the population average. Subsequent analyses have shown that this optimal strategy is located in a fragile region of the dosing space, suggesting that other treatment regimens would lead to more robust results in heterogenous cohorts (Jenner et al. 2021b).

Thus, VCTs have emerged as platforms with which to interrogate heterogeneous responses to drugs, delineate effective scheduling, and improve drug administration (Wang et al. 2008; Barish et al. 2017; Pérez-García et al. 2019; Cassidy and Craig 2019). In preclinical research, VCTs can also contribute to the decision-making process by distinguishing drug regimens leading to therapeutic successes and failures (Allen et al. 2016; Alfonso et al. 2020), and help to decipher individual patient risk classes and optimize drug-specific parameters (Viceconti et al. 2016; Boem et al. 2020). Across the multiple applications of virtual clinical trial strategies, the generation of virtual patient populations is paramount. Unfortunately, there is no solid consensus on the means of generating virtual patients. Here we address this specific issue by outlining popular approaches to virtual patient generation and highlighting their advantages and disadvantages using three case studies.

### USING POPULATION PHARMACOKINETIC MODELS TO GENERATE PATIENTS

In pharmacometric analysis, the standard method of evaluating and predicting the kinetics of plasma drug concentrations is through the assessment of drug pharmacokinetics (“what the body does to the drug”). Population pharmacokinetic (PopPK) models are built to discern population- and individual-level PKs using non-linear mixed effects (NLME) modelling. NLME models are statistical models that assume a fixed effect for the population and represent individual variation in the form of random effects. Let *P* be a set of parameters in the PK model. These parameters typically include factors like bioavailability (*F*), volume of distribution (*V*_d_), transit rates (*k*_*ij*_, where *i* and *j* denote model compartments), and clearance (*CL*). In its simplest form, a PopPK model for an individual *k* is given by the parameter vector *P*_*k*_ calculated as

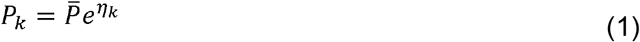

where 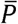 is the set of fixed (or population-level) parameters, *η*_*k*_ is a normally-distributed random variable with mean of 0 and variance 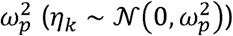, and 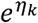 represents the resulting individual variability (IIV). Building a PopPK model using NLME models relies on population-level empirical data (usually from a clinical trial) from which the best pharmacokinetic model is defined. Here the “best” model is heuristic and determined by evaluating the calculated objective function after fitting several PK models. Multiple software packages, including R, NONMEM, and Monolix, can be used to perform NLME estimates and establish population PK models from data.

As PopPK models are empirically established and not generally built from mechanistic principles, they are primarily relevant for the specific population upon which they were constructed. Nonetheless, during drug development, we may be interested in establishing dosing strategies, exploring the potential for toxicity due to reduced kidney or liver function, etc. within a given population that go beyond the scenarios explored in a clinical trial. Here, leveraging a PopPK model is particularly attractive because it inherently accounts for individual variation within the population studied. To demonstrate how a PopPK model can be used to generate a virtual cohort, we considered a simple model of Gompertzian tumour growth and its treatment by gemcitabine, a synthetic pyrimidine nucleoside prodrug used as a chemotherapeutic agent in a variety of solid tumours (Joerger et al. 2014).

Let *N*(*t*) be the number of tumour cells at time *t*. The Gompertz model given by

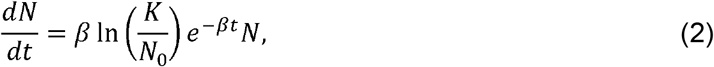

models saturable sigmoidal tumour growth to a carrying capacity of *k*. Here, *β* denotes the tumour growth rate (that decreases exponentially in time), and *N*_0_ the initial number of tumour cells. The PK model of gemcitabine is given by

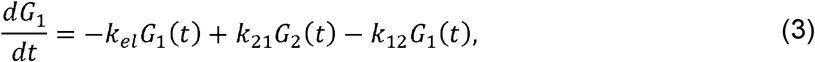

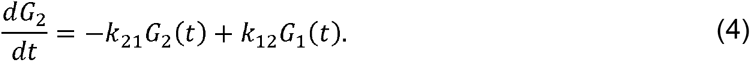

Each equation represents one of two compartments defined through a previous PopPK analysis (Jiang et al. 2007). These equations model the transfer of the drug between the central (*G*_1_(*t*)) and peripheral (*G*_2_(*t*)) compartments at rates *k*_12_ and *k*_21_, respectively. Gemcitabine was modelled as being administered directly into the central compartment from which it is eliminated linearly at rate (*k*_*el*_):

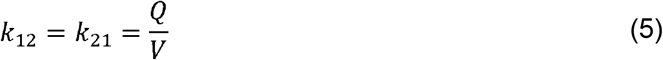

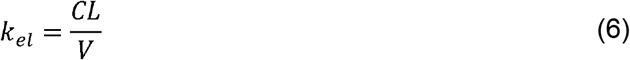

The between subject variability (BSV%) for *Q,CL,V* and the range for body surface area (*BSA*) were previously estimated using NLME modelling (Jiang et al. 2007). Here, BSA is used to calibrate gemcitabine doses.

We modelled the cytotoxic effects of chemotherapy on tumour growth using an inhibitory Hill effect function given by

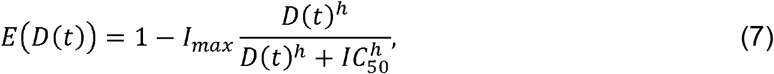

where *E*(*D*(*t*)) denotes the effect of a drug *D*(*t*) at time *t, I*_*max*_ represents the drug’s maximum inhibitory effect, *IC*_50_ the drug concentration at which inhibition is 50% its maximum, and *h* is the usual Hill coefficient controlling the slope of the curve. Integrating this into Eq. 2 we have the following model for the effects of gemcitabine on tumour growth

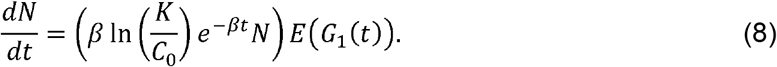

To generate virtual patients, we considered tumour cell parameters to be fixed at their previously estimated values and varied only the PK parameters according to the PopPK model for gemcitabine (**Table *1***). Assuming BSA to be normally distributed (Sacco et al. 2010), we calculated the mean and standard deviation required for the generation of BSA values (Hozo et al. 2005).

**Table 1.** Parameter values for gemcitabine inhibition of tumour growth. Parameters with reported variation were used to generate the virtual cohort.

| Parameter | Units | Value | BSV% | Ref |
| --- | --- | --- | --- | --- |
| $\beta$ | day<sup>-1</sup> | 0.0251 | - | Chosen |
| $K$ | cells | $5.157 \times 10^8$ | - | Chosen |
| $C_0$ | cells | $2 \times 10^6$ | - | Chosen |
| $I_{max}$ | unitless | 0.768 | - | Chosen |
| $h$ | unitless | 0.518 | - | Chosen |
| $IC_{50}$ | $\mu M$ | 297 | - | Chosen |
| $Q$ | L/min | 0.7 | 44 | (Jiang et al. 2007) |
| $CL$ | L/min | 2.7 | 31 | (Jiang et al. 2007) |
| $V$ | L | 15 | 39 | (Jiang et al. 2007) |
| $BSA$ | m<sup>2</sup> | Median: 1.8 | Range: 1.2 – 2.5 | (Jiang et al. 2007) |

We then sampled a set of parameter values using between-subject variability specific to that parameter. For example, a vector of normally distributed values with mean (e.g. 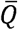) and standard deviation based on the BSV% of *Q* was used to generate a set of values {*Q*_*k*_}, *k* ∈ 1,2,3, …, *n* (where *n* is the cohort size) using

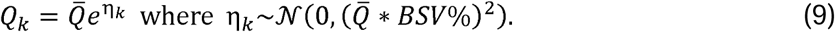

The *j*^th^ virtual patient is then given by the *j*^th^ value from the PK vector such that each patient is described as a set *P*_*j*_ = (*Q*_*j*_,*CL*_*j*_,*V*_*j*_,*BSA*_*j*_). Transit rates (*k*_12_,_*j*_ and *k*_21,*j*_) and elimination rates (*k*_*el,j*_) for each patient *j* were calculated using Eqs. 5 and 6, and the dose for each patient *j* was calculated using the dosage 1000mg/m^2^, where *BSA*_*j*_ (m^2^) is patient-specific body surface area. Using this process, we generated 200 virtual patients with their own dosage and model parameters tied to their assigned PK values.

A schematic of the PopPK/PD model is provided in **Figure 2A**. As expected, model simulations predicted that dynamics of cancer cell growth in the virtual cohort differed based on individual patients’ PK parameters (**Figure 2**). For the most part, we observed that drug concentrations fell within expected ranges in the central gemcitabine compartment (**Figure** 2) and peripheral compartment (**Figure *2***). The pharmacokinetics of gemcitabine have been shown to be linear up to 2500 mg/m^2^ which coincides closely with maximum-tolerated dose in a dose escalation study (Fossella et al. 1997). The dynamic of such a high gemcitabine initial concentration can be seen in red in **Figure *2*C-F**. However, the virtual population contained obvious outliers (**Figure *2***), with more than one virtual patient exhibiting markedly different drug concentrations from the rest of the cohort to the extent where toxic or lethal concentrations of gemcitabine were predicted. This highlights an obvious downfall of using PopPK models without integration of prior knowledge of inter-parameter relationships to generate virtual patients. Since the “top-down” approach (versus the “bottom-up” of mechanistic models) used here is not constrained by known mechanistic interactions, unrealistic (potentially dangerous) outcomes can be generated from what we may believe to be reasonable parameter ranges. This issue arises due to the method used to randomly sample parameter values from the previously established PopPK model with no built-in approach to verify that any specific combination of parameters (i.e. a virtual patient) is physiologically realistic. Though we may be able to clearly distinguish virtual patient outliers visually and remove them from the cohort, a systematic approach to remove parameter sets that generate unrealistic individuals despite being drawn from realistic distributions is needed.

**Figure 1.**
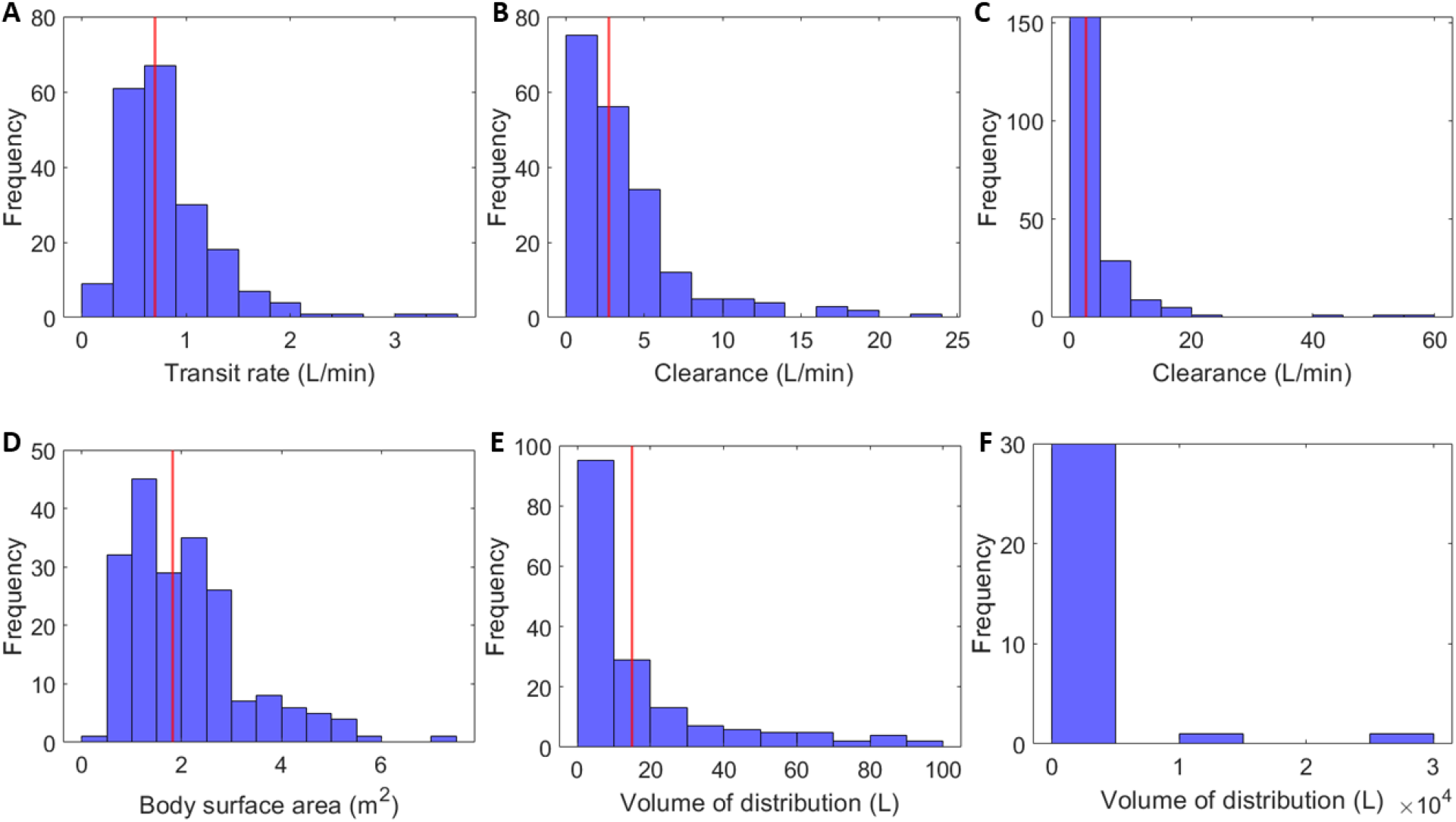
Histograms of parameter values generated for 200 virtual patients. A) Q, B) CL under 30L/min, C) CL including outliers, D) BSA, E) V under 100L and F) V outliers above 100L. The red line on each plot represents the average or median value found in **Table 1**.

**Figure 2.**
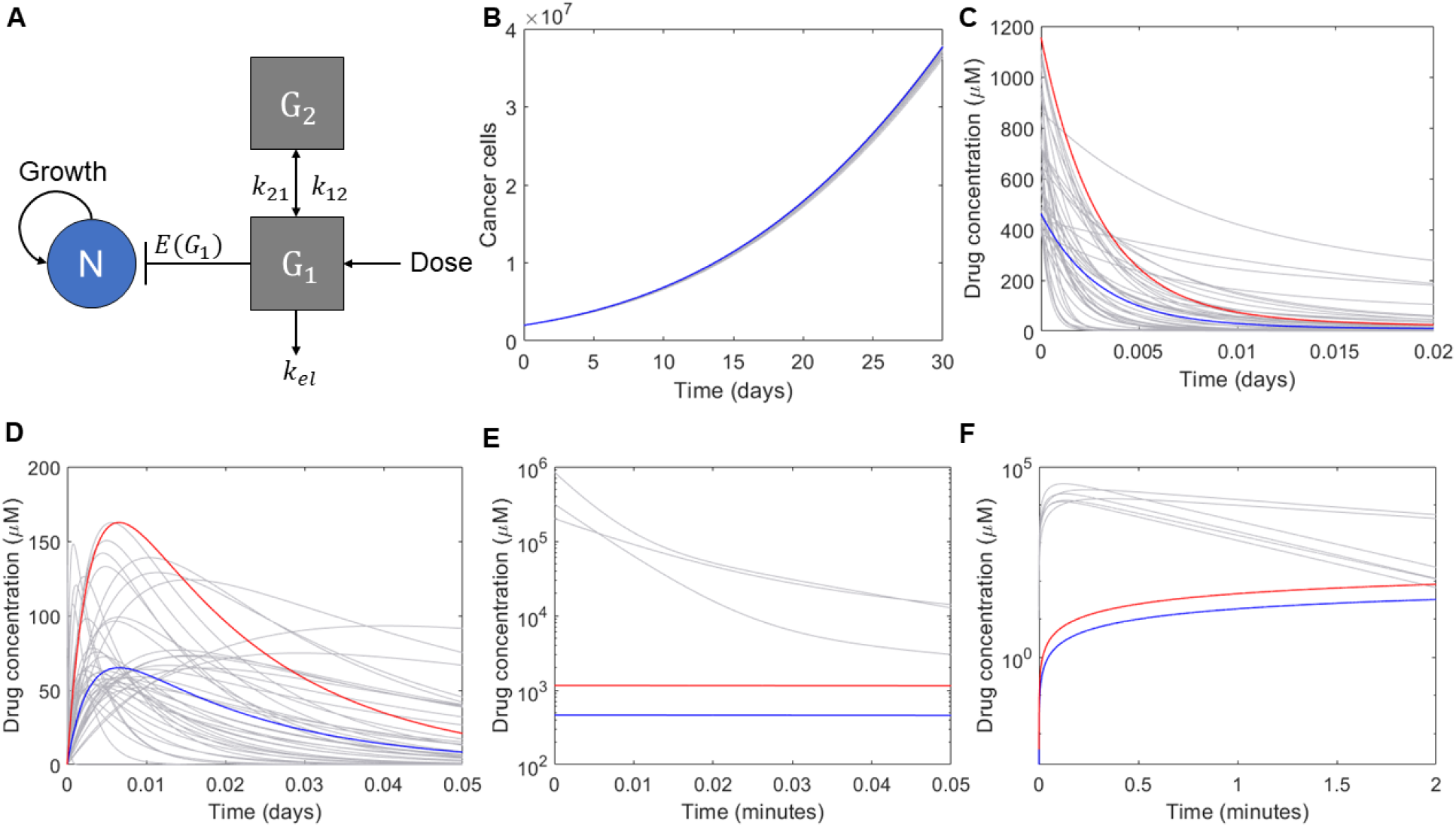
Tumour growth and gemcitabine concentrations in a cohort of 200 virtual patients. A) Schematic of the model and dynamics of B) cancer cell growth (*N*(*t*)) over 30 days. C) The central gemcitabine compartment (*G*_1_(*t*)) and D) the peripheral gemcitabine compartment (*G*_2_(*t*)) for reasonable drug doses and concentrations below a near maximum dose of 2500mg/m (red). Outliers in E) the central gemcitabine compartment (*G*_1_(*t*)) and F) the peripheral gemcitabine compartment (*G*_2_(*t*))compared to the average PopPK patient following different initial doses. Blue solid lines: average cohort response after 1000 mg/m^2^ dose of gemcitabine; red solid lines: average cohort response after 2500mg/m^2^ dose of gemcitabine; grey solid lines: individual virtual patients.

We can better understand how this random mismatch of parameters leads to potentially problematic virtual patients and outcomes by analyzing the PK parameters we generated and their implementation within the model considered here. As mentioned above, the elimination rate *k*_*el,j*_ for each patient was calculated by dividing their clearance (*CL*_,*j*_) by their compartment volume (*V*_,*j*_) (Eq. 6). Theoretically, because all parameter values are randomly drawn from established ranges, it is possible that a high clearance and a small volume may be paired. However, at the extreme for each parameter, this combination is likely to be unrealistic. This problem is further compounded as dose sizes are calculated for each patient based on their BSA, and empirically-estimated correlates of BSA to the other model parameters were not reported. For example, it would be rare to have a patient with a lower BSA relative to the cohort mean, but with higher compartment volumes and low transit and thus elimination rates. The lack of explicit physiological constraints during the virtual patient generation phase provides no explicit guarantee that the random combinations of parameters will lead to realistic individuals.

Despite these shortcomings, the generation of virtual patients from PopPK models is a simple process that can be implemented rapidly and with ease. Therefore, to mitigate the risks of generating non-viable virtual individuals from PopPK models, we suggest a thorough investigation of the parameter combinations resulting from the sampling process. Furthermore, imposing limits on parameter ranges will help to ensure physiologically relevant sampling is acheived. This process should be guided by known physiology to not overly restrict sampling and virtual-patient generation, and thus reduce bias in the VCT.

### ESTABLISHING THEORETICAL BOUNDS FROM EXPERIMENTAL MEASUREMENTS

In the previous section, we explored the generation of virtual individuals using only PK parameters while physiological parameters remained fixed. The emergence of quantitative system pharmacology (QSP) approaches has increasingly integrated detailed mechanistic models of physiological systems and disease processes to PK/PD models to provide a more holistic understanding of the effects of xenobiotics. We have previously shown that QSP models account for interindividual PK variability by the nature of their construction (Craig et al. 2016; Le Sauteur-Robitaille et al. 2021). Therefore, it is reasonable to generate virtual patients by uniquely varying physiological parameters in the model, since it is primarily physiological heterogeneity driving variable responses to drugs.

To demonstrate the generation and use of a virtual clinical trial incorporating virtual patients generated by sampling physiological parameters from theoretically-defined parameter ranges, we considered a model for the interaction between a cytotoxic chemotherapy drug ([*drug*] ng ·ml^-1^), a population of tumour cells (*S*(*t*)), and the immune system (*T*(*t*)). In this model, we investigated how the introduction of a chemotherapy drug may impact the antitumour immune response by reducing the pool of tumour cells and hence affecting the immune recruitment. We model cancer cells as growing logistically with proliferation rate *r* (day^-1^) and carrying capacity *K* (cells). The effects of the immune system on the tumour are modelled by supposing that cancer cells undergo apoptosis through contact with immune cells at a rate *k* (cells^-1^ day^-2^). We assumed that immune cells are recruited at a rate proportional to the amount of tumour cells *a* (day^-1^) and die at a rate *d* (day^-1^), giving the model

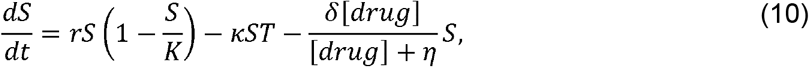

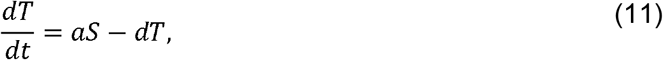

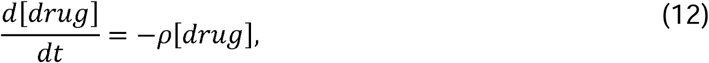

A schematic summary of the system can be found in **Figure 3A**.

**Figure 3.**
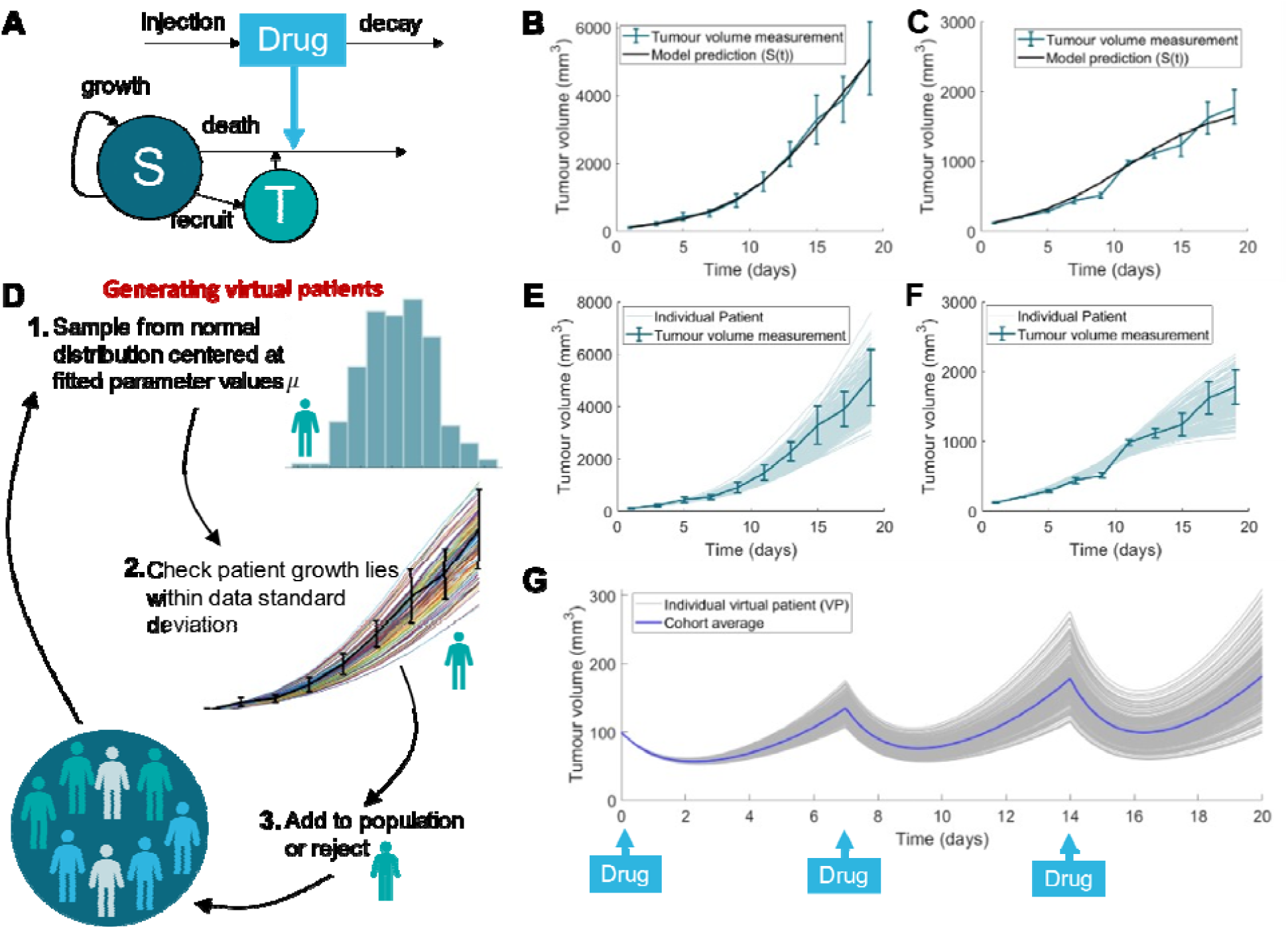
Virtual cohort example in immune-tumour-anti-cancer drug model. A) Model schematic. Cancer cells grow logistically and are killed by a cytotoxic chemotherapy drug ([ ]) and through tumour-immune () interactions. B)-C) Resulting model trajectories after parameter fits to untreated tumour growth without immune (B, control data (Oh et al. 2017)) and untreated tumour growth in the presence of the immune system (C, immune-suppressed data (Kim et al. 2011)). D) Schematic overview of the generation of virtual patients informed by experimental data. E-F) Predicted responses of tumour volume under control and treatment scenarios of the virtual patients accepted into the virtual patient cohort (light blue) with corresponding data measurements. G) Evolution of virtual cohorts’ tumour volume under chemotherapy treatment where the drug was administered every 7 days. Individual patient predictions (grey) and the cohort average evolution (blue) is given.

The model was parameterized using data of tumour growth in the absence of the immune system (Oh et al. 2017) (control tumour growth) and data of tumour growth in the presence of the immune system, but under suppressive virotherapy (Kim et al. 2011). We assumed the latter data to be representative of the immune-suppressed tumour growth model. Tumour volume was measured using callipers in mice and the average was used to fit parameters in the model. The data here were deployed solely for illustrative purposes, and we parameterized the model to the data using a simultaneous fitting approach (Gray and Coster 2016; Jenner et al. 2018).

Simultaneous fitting to the data was performed by setting the appropriate parts of the model to zero for each data set. In other words, for the control data, the drug and the immune population were set to zero and for the suppressed tumour growth data (immune-present data) the drug parameters and variables were set to zero. The remaining parameters in the drug-free model, *r,K,k,a* and *d* were then fit simultaneously using non-linear least squares fitting with the objective of minimizing the residual of the model to both sets of data simultaneously. The fitted parameters can be found in **Table 2**, and the resulting model approximation to the data is in **Figure 3B** and **3C**. To parametrize the models for chemotherapy drug decay and drug-induced tumour cell death we used parameters for PAC-1 (first procaspase activating compound) fit previously from *in vitro* experiments (Crosley et al. 2021).

**Table 2.** Parameter values for the tumour growth under immune suppression and chemotherapy treatment. Parameters for the tumour growth and immune dynamics were obtained through fitting to data (Kim et al. 2011; Oh et al. 2017). The remaining parameters were estimated from PAC-1 (first procaspase activating compound) dynamics (Crosley et al. 2021).

| Parameter | Units | Description | Value | Ref |
| --- | --- | --- | --- | --- |
| $r$ | $\text{day}^{-1}$ | Tumour cell proliferation rate | 0.2349 | Fit **Fig. 2** |
| $K$ | cells | Tumour carrying capacity | 10190 | Fit **Fig. 2** |
| $\kappa$ | $\text{day}^{-1}\text{cells}^{-1}$ | Immune cell induced tumour cell death rate | 0.0021 | Fit **Fig. 2** |
| $a$ | $\text{day}^{-1}$ | Immune cell recruitment rate | 7.086 | Fit **Fig. 2** |
| $d$ | $\text{day}^{-1}$ | Immune cell death rate | 162.5 | Fit **Fig. 2** |
| $\delta$ | $\text{day}^{-1}$ | Drug induced tumour cell death | 2.64 | Crosley et al. 2021 |
| $\eta$ | $\text{ng}\cdot\text{ml}^{-1}$ | Drug half-effect concentration | 5 | Crosley et al. 2021 |
| $\rho$ | $\text{day}^{-1}$ | Drug elimination rate | 0.903 | Crosley et al. 2021 |

Next, to create *n* virtual patients, we sampled parameters from a normal distribution centered at *μ* with standard deviation *σ*, rejecting any negative parameters (**Figure 3D**). We fixed ***μ*** to be the set of parameter values obtained from our fitting procedure for *r,K,k,a* and d as we assumed individuals would vary in both the tumour growth and immune dynamics. The standard deviation ***σ*** was fixed to ***σ*** = 0.2***μ*** to minimize the likelihood of sampling negative parameter values while maximising the variation on the parameters chosen for our virtual population. Sampling with this ***σ*** resulted in 634 random samples being rejected to make 200 virtual patients. Increasing to ***σ*** *=* 0.5***μ*** resulted in a higher number of negative parameter samples, which are set to zero, and also a higher number of samples rejected (1917). It is possible to also estimate ***σ*** from the standard deviation in the data (Jenner et al. 2021a).

To create a realistic representation of patients, we restricted the inclusion of virtual patients to those whose simulated control tumour growth and immune-suppressed tumour growth was within physiological reasonable regimes which we designated to be three standard deviations 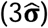 of the mean control data and five standard deviations 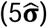 of the immune suppressed tumour growth data (**Figure 3B** and **3C**). Different thresholds where chosen for each case as it was not possible to generate patients within 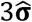 of tumour measurements at day 9 and 10 in the immune-suppressed data (Figure 3F) with a fixed underlying parameter distribution for sampling. As such, we increased the tumour volume trajectories for the immune suppressed data to lie within 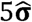 of the data. Any virtual patient whose simulated tumour growth in the absence of drug lay outside these intervals was rejected from the virtual trial. Remaining patients were accepted into the trial and the distribution of all model-predicted tumour growths lay close to the data (**Figure 3E** and **3F**). This step mimics the selection process of clinical trials (for example, including only those patients with a specific grade of a cancer).

Following this generation approach, tumour growth in both the control and treatment scenarios was confirmed to fall within the prescribed heterogeneity bounds (**Figure 3E** and **3F**). Over time, we see the variation in virtual patient growths increasing with large variations of patient dynamics by day 19 in both the control and immune-suppressed case. Simulating drug administration every seven days starting at day 0, we observe a similar trend, where virtual patients respond similarly in the first cycle before more significant differences in tumour volume emerge (**Figure 3G**).

As in the approach described in the previous section, the techniques described here are relatively straightforward to implement. However, in contrast to basing the virtual patient generation solely on PopPK parameters, this technique integrates constraints on the mechanistic (or physiological) model parameters through the integration of biological experimental data. Unfortunately, the procedure described above is still not able to capture parameter correlations (for example, a patient’s tumour growth rate (*r*) and carrying capacity (*K*)), although it would be possible to address this shortcoming by performing statistical tests to guide the parameter sampling. The additional step of comparing model prediction to measured outputs responds to the limitation of creating unrealistic virtual patients. In practice, a large number of proposed virtual patients may need to be generated in order to ensure that the final trial cohort is sufficiently large In this section, virtual patients trajectories were restricted to lying between 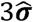 and 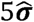 for the control and immune-suppressed data respectively. The difference in these ranges for the physiological regimes was driven by the inability to achieve trajectories within 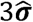 of the immune-suppressed data. As described in the next section, this limitation can be overcome by adding a step within the virtual patient generation phase to ensure that all parameter samplings result in model trajectories that describe clinical observations.

### QUANTITATIVE SYSTEMS PHARMACOLOGY APPROACHES

As more mechanisms are included, model complexity and the sparsity of relevant data complicates the implementation of VCTs. Accordingly, methods to generate virtual populations that reproduce the heterogeneity in patients as well as allow for the exploration of parametric uncertainty have been devised to overcome these challenges. One of the best-known approaches in this vein is that of Allen et al. 2016 who proposed a method to generate a large cohort of heterogeneous virtual patients by sampling a parameter set from a bounded interval informed by physiological constraints, and optimizing predicted trajectories to ensure model outcomes are within clinically-observed ranges. Therefore, this methodology expands upon the approach studied in the previous section, by explicitly integrating the constraint that model predictions for each virtual patient must fall within empirically-determined ranges, into the virtual patient generation process through an optimization step.

To demonstrate the generation of a virtual patient population using this approach, we will again use the cancer-immune interaction model introduced in Eqs. 10-12. Here we limit our focus solely on generating virtual patients using this alternative approach and thus ignore the effect of the drug, and reduce the model to a system of ODEs describing the time evolution of tumour (*S*(*t*)) and immune (*T*(*t*)) cells:

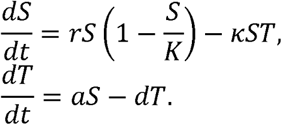

All parameters are as previously defined in **Table 2**. A schematic of various steps involved in virtual patient generation using the Allen et al. 2016 method is presented in **Figure 4**.

**Figure 4.**
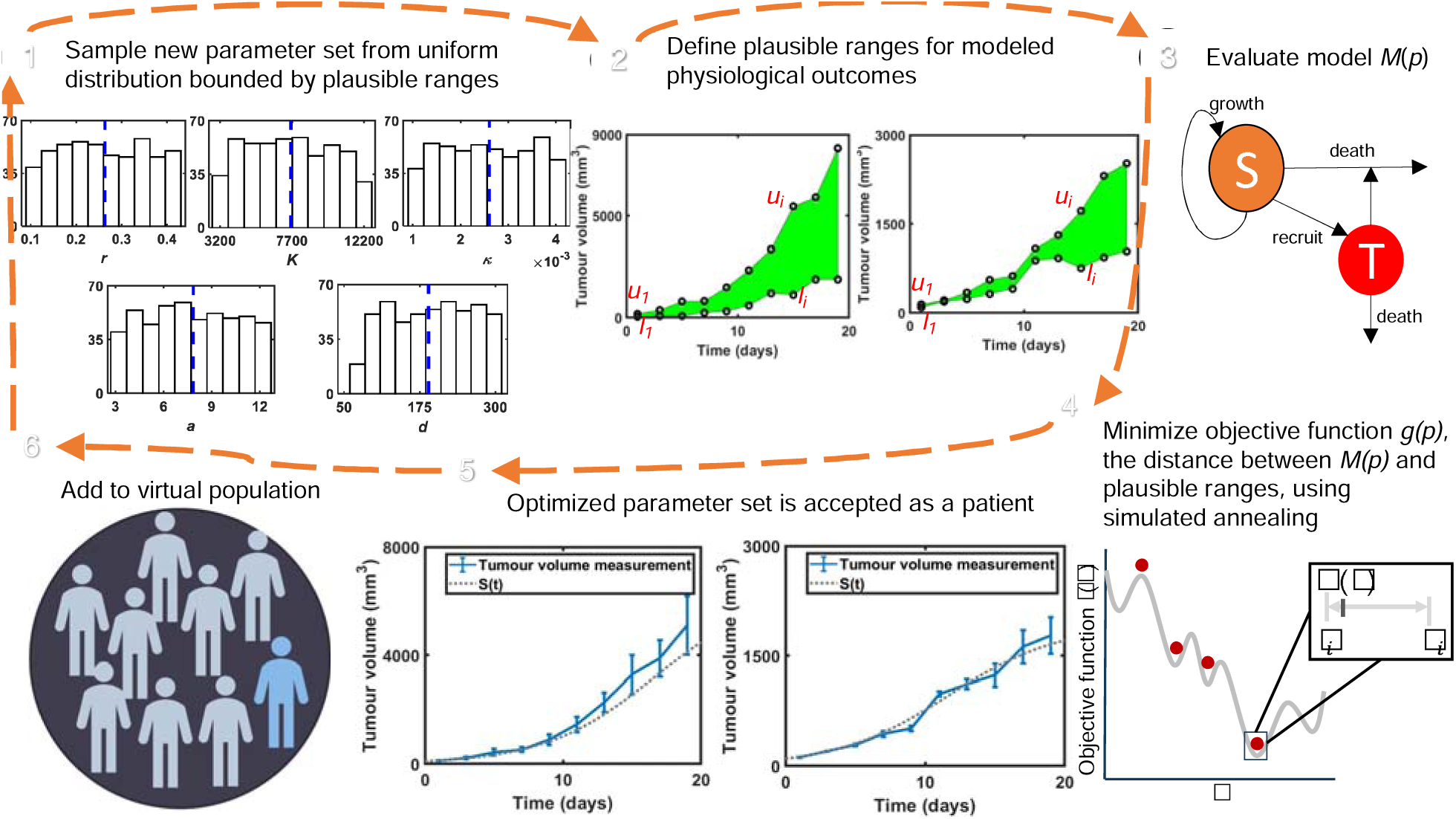
Major steps in virtual patient generation using the Allen et al. method. **1)** Initial model parameters are sampled from a uniform distribution bounded by plausible ranges. **2)** Define an upper (*u*_i_) and lower (*l*_i_) bound for the observable model outcome (tumour volume) at each time point of the available data. **3)** The model is evaluated with the initial parameter guesses sampled from the uniform distribution and obtain model output, *M*(*p, t*). **4)** An optimal parameter set that minimizes the objective function *g(p)* is obtained using the simulated annealing algorithm. **5)** The optimization of *g(p)* using the simulated annealing algorithm generates a virtual patient whose disease dynamics lie within the physiologically plausible ranges. **6)** The newly generated patent is added to the virtual patient population. The optimization routine is repeated until the required number of patients are generated.

A key component that determines the successful implementation of the Allen et al. 2016 method is the ability to define realistic bounds for the model parameters. In typical applications, these bounds can be inferred through empirical estimates from physiological experiments or through theoretical considerations. In our simulations, we consider parameter values within three standard deviations (3 ***σ***) from mean (***μ***) values presented in **Table 2** as plausible ranges for the model parameters, *r, K, k, a* and *d*. As we described in the previous section, the mean parameter values are obtained by non-linear least-squares fitting. In the prior section, each parameter value was assumed to be normally distributed with standard deviation ***σ*** = 0.2***μ***, so here we are considering a much broader range of plausible parameter values.

To start the patient generation process, we constructed an initial parameter set where each parameter is drawn from a uniform distribution bounded by the plausible ranges discussed above. Next, we optimized each set of parameters (*r, K, k, a* and *d*) using simulated annealing (a probabilistic optimization algorithm implemented as the *simulannealbnd* function in MATLAB (Mathworks 2020) to ensure that predicted model trajectories fall within physiological ranges. For our purposes, these bounds on model outputs were assumed to be within three standard deviations of the control and immunosuppressed tumour growth data means. We defined the objective function of the simulated annealing scheme to be

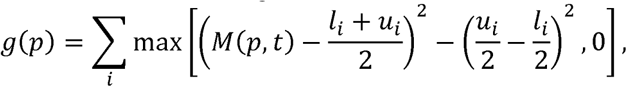

and minimized *g(p)*. Here, *M* (*p, t*)denotes the model output for parameter set *p* at time *t* and *l*_*i*_ and *u*_*i*_ denotes the *i*^th^ plausible upper and lower bounds of the data. By defining the objective function in this fashion, we are guaranteed that the contribution of *M*(*p, t*) − (*l*_*i*_ + *u*_*i*_)/2 is zero if the parameters lead to an outcome within the plausible range. If the optimization converges, the resulting parameter set was considered to belong to a physiologically-valid virtual patient and was added to the trial population.

Following these steps, we generated a virtual population of 500 patients using the Allen et al. 2016 method. The trajectories of all virtual patients were confirmed to lie within the physiological bounds determined for the model outputs (**Figure 5C** and **Figure 5D**). Thus, this method generates a large pool of patients with realistic disease dynamics without manual verification as was required in the methods discussed in Sections 2 and 3. It should be noted that for specific applications, e.g. to consider patients with certain levels of tumour growth, we can subsample from the resulting virtual patient cohort. We may also exploit predetermined, empirical parameter distributions to create specific subpopulations of patients. For example, the posterior distribution of both *r* and *k* were found to differ significantly from their prior (**Figure 5A** and **5B**), suggesting a mechanistic role of both parameters that may drive heterogeneity and segregate patient responses. Narrowing in on specific values of each (in isolation and in combination) could reveal specific subpopulations of virtual patients and allow for further tailoring of treatment strategies.

**Figure 5.**
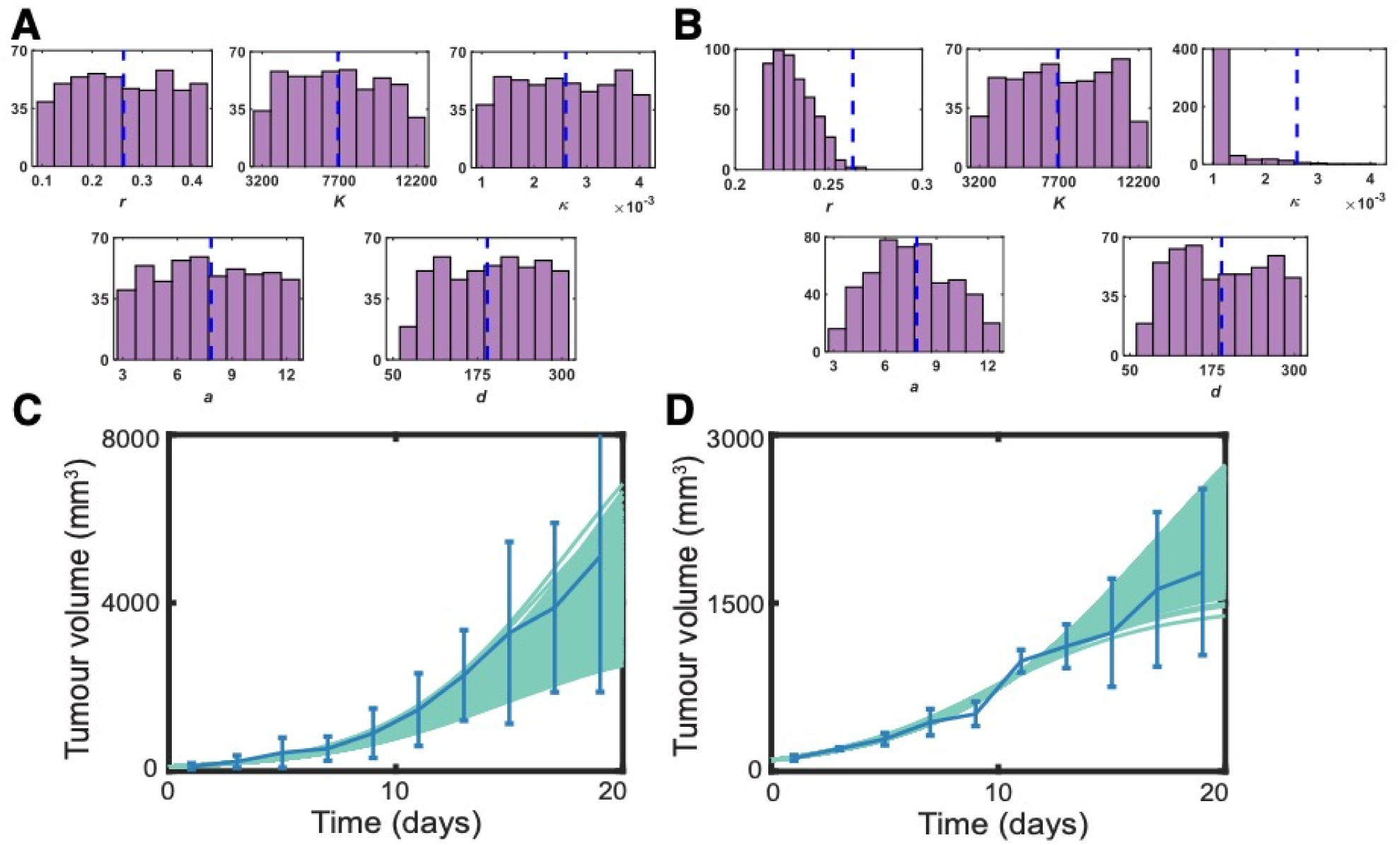
A virtual population of 500 patients generated using the Allen et al. method. **A)** shows the initial parameter value distributions (uniformly distributed) and **B)** shows the final parameter value distributions after the optimization using the simulated annealing algorithm. The blue vertical line on each histogram represents the mean value of the distribution. **C)** and **D)** show the tumour growth in the control and in presence of the immune system, respectively. The virtual patient dynamics trajectories (cyan) are superimposed with the corresponding data measurements (blue). The error bars denote three standard deviations from the data mean.

We have seen that the generation of VPs from PopPK models may lead to patients with disease dynamics beyond physiological limits, when using random parameter sampling without any built-in mechanism to incorporate physiological constraints and correlations. In contrast, the Allen et al. approach, which leverages physiologically-informed bounds on parameters and model outputs together with robust optimization using the simulated annealing algorithm, ensures the generation of realistic patient cohorts. While the second method we described based on obtaining bounds from experimental measurements is a reliable approach to get a realistic patient population with dynamics within the physiological bounds, the brute force way of generating a large number of candidate patients to eventually select the total size of the cohort is not efficient. On the other hand, the Allen et al. 2016 method is an alternative that produces a patient cohort without relying on generating a large candidate population. This comes, however, at a computational cost as every virtual patient must undergo optimization prior to being accepted into the virtual cohort. In response, recent extensions of the method involving augmenting optimization of the cost function (nested simulated annealing) or adopting alternative optimization routines (modified Metropolis-Hastings and genetic algorithms) have been shown to lead to more efficient generation of virtual patients (Rieger et al. 2018).

## DISCUSSION

Designing and developing new xenobiotics in immuno-oncology is complicated and costly, driven by our sometimes-limited knowledge of the mechanisms regulating therapeutic efficacy. Mathematical and computational modelling are increasingly integrated along the drug development pipeline and in fundamental studies to help identify regulators of drug responses. Given the high degree of variability observed within patient populations, quantitative approaches that can also capture the heterogeneity in outcomes are now frequently deployed to assess the degree to which variability affects outcomes and to discern the sources of such heterogeneity.

*In silico* clinical trials are therefore well-situated to help guide the preclinical-to-clinical translation of candidate molecules, to assess the best candidate populations for a given treatment, to delineate successful combination strategies, and to establish optimal dosing schedules. Virtual patient populations are at the heart of these computational trials and must be reflective of the variability we observe in real patient populations. However, there is no universally accepted method of generating a virtual patient population, and each approach comes with its own set of advantages and limitations. Here we have described three widely-deployed methods to generate a virtual cohort, starting from the most simple implementation using empirically-defined population pharmacokinetic models to the more complex, and perhaps most robust, approach described by Allen et al. 2016 that verifies a virtual patient’s trajectory within the generation step.

Whether the patient population is generated using statistical methods or probabilistic data-fitting methods, it is clear that the choice of which parameters are used to define the population, and thus which remain constant across the population, depends on the mathematical model and the application. For models with a large number of model parameters, a high-dimensional patient-specific parameter set may be challenging to both appropriately set-up and to optimize. Consideration of key aspects such as model identifiability, parameter sensitivity, and significant biological mechanisms can help inform which parameters to include in the patient-specific set. Additionally, the assumptions made during the set-up phase of VCT construction will have implications on the conclusions of the simulated study, and these implications should be explored. Further, as new methods for creating virtual patients continue to be proposed in the literature, it is important to establish standards for virtual clinical trials, in terms of the data needed to perform them, the dimensionality of the patient-specific parameter sets, how interpatient variability is represented by the data, and how the output of these trials is used and validated. Clearly there is still lots of work to be done in the theory of VCT design, implementation, and assessment. To that end, new approaches integrating machine learning and data dimensionality reduction techniques may also prove useful for selecting parameters of interest to generate the virtual cohort.

Ultimately, the choice of method for generating a virtual cohort depends on a number of factors, most predominately the level of complexity required, and the time allowed for model generation and implementation of the method. Once created, virtual patient populations and *in silico* clinical trials are powerful new tools that can provide biological insights that may be difficult or impossible to otherwise identify (Jenner et al. 2021a). When used in complement to experimental and clinical studies, virtual clinical trials have the potential to markedly decrease attrition rates along the drug development pipeline, helping to reduce disappointing trial results and improve patient outcomes.

## ACKNOWLEDGEMENTS

AS was funded by a CRM-ISM Postdoctoral Fellowship, JLR received funding from a CAMBAM fellowship and ISM scholarship, and an NSERC Discovery Grant to MC. KPW was supported by an NSERC Discovery Grant RGPIN-2018-04205. MC was funded by NSERC Discovery Grant RGPIN-2018-04546, and an FRQS J1 Research Scholar Grant. Some of the work in this chapter came from the tutorials that were run during the Centre de recherches mathématiques and Centre for Modelling in the Biosciences and Medicine funded workshop *Computational modelling of Cancer Biology and Treatments* supporting ALJ and MC.

## CONFLICT OF INTEREST DISCLOSURE

The authors declare that no conflicts of interest exist.

## Notes

### Competing Interest Statement

The authors have declared no competing interest.

## REFERECES

Agosti A, Giverso C, Faggiano E, et al (2018) A personalized mathematical tool for neuro-oncology: A clinical case study. Int J Non Linear Mech 107:170–181. https://doi.org/10.1016/j.ijnonlinmec.2018.06.004

Alfonso S, Jenner AL, Craig M (2020) Translational approaches to treating dynamical diseases through in silico clinical trials. Chaos An Interdiscip J Nonlinear Sci 30:123128. https://doi.org/10.1063/5.0019556

Allen RJ, Rieger TR, Musante CJ (2016) Efficient Generation and Selection of Virtual Populations in Quantitative Systems Pharmacology Models. CPT Pharmacometrics Syst Pharmacol 5:140–146. https://doi.org/10.1002/psp4.12063

An G (2004) In silico experiments of existing and hypothetical cytokine-directed clinical trials using agent-based modeling*. Crit Care Med 32:2050–2060. https://doi.org/10.1097/01.CCM.0000139707.13729.7D

Andtbacka RHI, Ross M, Puzanov I, et al (2016) Patterns of clinical response with talimogene laherparepvec (T-VEC) in patients with melanoma treated in the OPTiM phase III clinical trial. Ann Surg Oncol 23:4169–4177

Andtbacka Rhii, Kaufman HL, Collichio F, et al (2015) Talimogene laherparepvec improves durable response rate in patients with advanced melanoma. J Clin Oncol 33:2780–2788. https://doi.org/10.1200/JCO.2014.58.3377

Barish S, Ochs MF, Sontag ED, Gevertz JL (2017) Evaluating optimal therapy robustness by virtual expansion of a sample population, with a case study in cancer immunotherapy. Proc Natl Acad Sci 114:E6277–E6286

Bland JM, Altman DG (2011) Comparisons against baseline within randomised groups are often used and can be highly misleading. Trials 12:264. https://doi.org/10.1186/1745-6215-12-264

Boem F, Malagrinò I, Bertolaso M (2020) In Silico Clinical Trials: A Possible Response to Complexity in Pharmacology. In: Uncertainty in Pharmacology. pp 135–152

Bozic I, Antal T, Ohtsuki H, et al (2010) Accumulation of driver and passenger mutations during tumor progression. Proc Natl Acad Sci 107:18545–18550. https://doi.org/10.1073/pnas.1010978107

Brown BW (1980) The Crossover Experiment for Clinical Trials. Biometrics 36:69. https://doi.org/10.2307/2530496

Cassidy T, Craig M (2019) Determinants of combination GM-CSF immunotherapy and oncolytic virotherapy success identified through in silico treatment personalization. PLOS Comput Biol 15:e1007495. https://doi.org/10.1371/journal.pcbi.1007495

Clermont G, Bartels J, Kumar R, et al (2004) In silico design of clinical trials: A method coming of age. Crit Care Med. https://doi.org/10.1097/01.CCM.0000142394.28791.C3

Corral-Acero J, Margara F, Marciniak M, et al (2020) The ‘Digital Twin’ to enable the vision of precision cardiology. Eur Heart J 41:4556–4564. https://doi.org/10.1093/eurheartj/ehaa159

Craig M, González-Sales M, Li J, Nekka F (2016) Impact of pharmacokinetic variability on a mechanistic physiological pharmacokinetic/pharmacodynamic model: a case study of neutrophil development, PM00104, and filgrastim. In: Toni B (ed) Mathematical Sciences with Multidisciplinary Applications. Springer Science + Business Media, New York, pp 91–112

Craig M, Kaveh K, Woosley A, et al (2019) Cooperative adaptation to therapy (CAT) confers resistance in heterogeneous non-small cell lung cancer. PLOS Comput Biol 15:1–19. https://doi.org/10.1371/journal.pcbi.1007278

Crosley P, Farkkila A, Jenner AL, et al (2021) Procaspase-Activating Compound-1 synergizes with TRAIL to induce apoptosis in established granulosa cell tumor cell line (KGN) and explanted patient granulosa cell tumor cells in vitro. Int J Mol Sci 22:4699. https://doi.org/10.3390/ijms22094699

DiMasi JA, Grabowski HG, Hansen RW (2016) Innovation in the pharmaceutical industry: New estimates of R&D costs. J Health Econ 47:20–33. https://doi.org/10.1016/j.jhealeco.2016.01.012

Emilia Kozlowska E, Färkkilä A, Vallius T, et al (2018) Mathematical modeling predicts response to chemotherapy and drug combinations in ovarian cancer. Cancer Resarch 78:4036–4044

Fossella F V, Lippman SM, Shin DM, et al (1997) Maximum-tolerated dose defined for single-agent gemcitabine: a phase I dose-escalation study in chemotherapy-naive patients with advanced non-small-cell lung cancer. J Clin Oncol 15:310–316. https://doi.org/10.1200/JCO.1997.15.1.310

Gray CW, Coster ACF (2016) The Akt switch model: Is location sufficient? J Theor Biol 398:103–111. \https://doi.org/10.1016/j.jtbi.2016.03.005

Gyuk P, Vassányi I, Kósa I (2019) Blood Glucose Level Prediction for Diabetics Based on Nutrition and Insulin Administration Logs Using Personalized Mathematical Models. J Healthc Eng 2019:1–12. https://doi.org/10.1155/2019/8605206

Hay M, Thomas DW, Craighead JL, et al (2014) Clinical development success rates for investigational drugs. Nat Biotechnol 32:40–51. https://doi.org/10.1038/nbt.2786

Hozo SP, Djulbegovic B, Hozo I (2005) Estimating the mean and variance from the median, range, and the size of a sample. BMC Med Res Methodol 5:1–10. https://doi.org/10.1186/1471-2288-5-13

Jafarnejad M, Gong C, Gabrielson E, et al (2019) A Computational Model of Neoadjuvant PD-1 Inhibition in Non-Small Cell Lung Cancer. AAPS J 21:79. https://doi.org/10.1208/s12248-019-0350-x

Jenner AL, Aogo RA, Alfonso S, et al (2021a) COVID-19 virtual patient cohort suggests immune mechanisms driving disease outcomes. PLOS Pathog 17:e1009753. https://doi.org/10.1371/journal.ppat.1009753

Jenner AL, Cassidy T, Belaid K, et al (2021b) In silico trials predict that combination strategies for enhancing vesicular stomatitis oncolytic virus are determined by tumor aggressivity. J Immunother Cancer 9:e001387. https://doi.org/10.1136/jitc-2020-001387

Jenner AL, Yun C-O, Kim PS, Coster ACF (2018) Mathematical modelling of the interaction between cancer cells and an oncolytic virus: insights into the effects of treatment protocols. Bull Math Biol 80:. https://doi.org/10.1007/s11538-018-0424-4

Jiang X, Galettis P, Links M, et al (2007) Population pharmacokinetics of gemcitabine and its metabolite in patients with cancer: effect of oxaliplatin and infusion rate. https://doi.org/10.1111/j.1365-2125.2007.03040.x

Joerger M, Huitema ADR, Koeberle D, et al (2014) Safety and pharmacology of gemcitabine and capecitabine in patients with advanced pancreatico-biliary cancer and hepatic dysfunction. Cancer Chemother Pharmacol 73:113–124. https://doi.org/10.1007/s00280-013-2327-2

Kim P-H, Sohn J-H, Choi J-W, et al (2011) Active targeting and safety profile of PEG-modified adenovirus conjugated with herceptin. Biomaterials 32:2314–2326. https://doi.org/10.1016/j.biomaterials.2010.10.031

Kirtane AR, Abouzid O, Minahan D, et al (2018) Development of an oral once-weekly drug delivery system for HIV antiretroviral therapy. Nat Commun 9:2. https://doi.org/10.1038/s41467-017-02294-6

Kozlowska E, Vallius T, Hynninen J, et al (2019) Virtual clinical trials identify effective combination therapies in ovarian cancer. Sci Rep 9:18678. https://doi.org/10.1038/s41598-019-55068-z

Le Sauteur-Robitaille J, Yu ZS, Craig M (2021) Impact of estrogen population pharmacokinetics on a qsp model of mammary stem cell differentiation into myoepithelial cells. AIMS Math 6:10861–10880. https://doi.org/10.3934/math.2021631

Lipsky MS, Sharp LK From idea to market: the drug approval process. J Am Board Fam Pract 14:362–7

Ma H, Pilvankar M, Wang J, et al (2021) Quantitative Systems Pharmacology Modeling of PBMC-Humanized Mouse to Facilitate Preclinical Immuno-oncology Drug Development. ACS Pharmacol Transl Sci 4:213–225. https://doi.org/10.1021/acsptsci.0c00178

Mathworks 2020 Matlab 2020b

Milberg O, Gong C, Jafarnejad M, et al (2019) A QSP Model for Predicting Clinical Responses to Monotherapy, Combination and Sequential Therapy Following CTLA-4, PD-1, and PD-L1 Checkpoint Blockade. Sci Rep 9:11286. https://doi.org/10.1038/s41598-019-47802-4

Mohs RC, Greig NH (2017) Drug discovery and development: role of basic biological research. Alzheimer’s Dement Transl Res Clin Interv 3:651–657. https://doi.org/10.1016/j.trci.2017.10.005

Oh E, Oh J-E, Hong J, et al (2017) Optimized biodegradable polymeric reservoir-mediated local and sustained co-delivery of dendritic cells and oncolytic adenovirus co-expressing IL-12 and GM-CSF for cancer immunotherapy. J Control Release 259:115–127. https://doi.org/10.1016/j.jconrel.2017.03.028

Pérez-García VM, Ayala-Hernández LE, Belmonte-Beitia J, et al (2019) Computational design of improved standardized chemotherapy protocols for grade II oligodendrogliomas. PLoS Comput Biol 15:1–17. https://doi.org/10.1371/journal.pcbi.1006778

Piantadosi S (2017) Clinical Trials: A Methodologic Perspective, 3rd edn. New York, New York, USA

Pitcher MJ, Bowness R, Dobson S, Gillespie SH (2018) A spatially heterogeneous network-based metapopulation software model applied to the simulation of a pulmonary tuberculosis infection. Appl Netw Sci 3:33. https://doi.org/10.1007/s41109-018-0091-2

Polasek TM, Rostami-Hodjegan A (2020) Virtual Twins: Understanding the Data Required for Model-Informed Precision Dosing. Clin Pharmacol Ther 107:742–745. https://doi.org/10.1002/cpt.1778

Rieger TR, Allen RJ, Bystricky L, et al (2018) Improving the generation and selection of virtual populations in quantitative systems pharmacology models. Prog Biophys Mol Biol 139:15–22. https://doi.org/10.1016/j.pbiomolbio.2018.06.002

Sacco JJ, Botten J, Macbeth F, et al (2010) The average body surface area of adult cancer patients in the UK: A multicentre retrospective study. LoS One 5:1–6. https://doi.org/10.1371/journal.pone.0008933

Sayama H, Marcantonio D, Nagashima T, et al (2021) Virtual clinical trial simulations for a novel KRAS G12C inhibitor (ASP2453) in non-small cell lung cancer. CPT Pharmacometrics Syst Pharmacol 10:864–877. https://doi.org/10.1002/psp4.12661

Seidel JA, Otsuka A, Kabashima K (2018) Anti-PD-1 and Anti-CTLA-4 Therapies in Cancer: Mechanisms of Action, Efficacy, and Limitations. Front Oncol 8:1–14. https://doi.org/10.3389/fonc.2018.00086

Sové RJ, Jafarnejad M, Zhao C, et al (2020) QSP-IO: A Quantitative Systems Pharmacology Toolbox for Mechanistic Multiscale Modeling for Immuno-Oncology Applications. CPT Pharmacometrics Syst Pharmacol 9:484–497. https://doi.org/10.1002/psp4.12546

Stadeli KM, Richman DD (2013) Rates of emergence of HIV drug resistance in resource-limited settings: A systematic review. Antiretrovir Ther 18:115–123

Switchenko JM, Heeke AL, Pan TC, Read WL (2019) The use of a predictive statistical model to make a virtual control arm for a clinical trial. PLoS One 14:e0221336. https://doi.org/10.1371/journal.pone.0221336

Tirosh I, Izar B, Prakadan SM, et al (2016) Dissecting the multicellular ecosystem of metastatic melanoma by single-cell RNA-seq. Sci (80-) 352:189–196. https://doi.org/10.1126/science.aad0501

Viceconti M, Henney A, Morley-Fletcher E (2016) In silico clinical trials: how computer simulation will transform the biomedical industry. Int J Clin Trials 3:37. https://doi.org/10.18203/2349-3259.ijct20161408

Visentin R, Dalla Man C, Kovatchev B, Cobelli C (2014) The University of Virginia/Padova Type 1 Diabetes Simulator Matches the Glucose Traces of a Clinical Trial. Diabetes Technol Ther 16:428–434. https://doi.org/10.1089/dia.2013.0377

Vodovotz Y, Billiar TR (2013) In Silico Modeling. Crit Care Med 41:2008–2014. https://doi.org/10.1097/CCM.0b013e31829a6eb4

Wang H, Ma H, Sové RJ, et al (2021) Quantitative systems pharmacology model predictions for efficacy of atezolizumab and nab-paclitaxel in triple-negative breast cancer. J Immunother Cancer 9:e002100. https://doi.org/10.1136/jitc-2020-002100

Wang H, Milberg O, Bartelink IH, et al (2020) In silico simulation of a clinical trial with anti-CTLA-4 and anti-PD-L1 immunotherapies in metastatic breast cancer using a systems pharmacology model. R Soc Open Sci 6:190366. https://doi.org/10.1098/rsos.190366

Wang Y, Bhattaram AV, Jadhav PR, et al (2008) Leveraging Prior Quantitative Knowledge to Guide Drug Development Decisions and Regulatory Science Recommendations: Impact of FDA Pharmacometrics During 2004-2006. J Clin Pharmacol 48:146–156. https://doi.org/10.1177/0091270007311111

Wilkie KP, Hahnfeldt P (2017) Modeling the Dichotomy of the Immune Response to Cancer: Cytotoxic Effects and Tumor-Promoting Inflammation. Bull Math Biol 79:1426–1448. https://doi.org/10.1007/s11538-017-0291-4

